# scSGL: Signed Graph Learning for Single-Cell Gene Regulatory Network Inference

**DOI:** 10.1101/2021.07.08.451697

**Authors:** Abdullah Karaaslanli, Satabdi Saha, Selin Aviyente, Tapabrata Maiti

**Affiliations:** Department of Electrical and Computer Engineering, Michigan State University, East Lansing, MI, USA; Department of Statistics and Probability, Michigan State University, East Lansing, MI, USA

**Keywords:** Graph Signal Processing, Signed Graph Learning, Gene Regulatory Network Inference

## Abstract

**Motivation:** Elucidating the topology of gene regulatory networks (GRNs) from large single-cell RNA sequencing (scRNAseq) datasets, while effectively capturing its inherent cell-cycle heterogeneity and dropouts, is currently one of the most pressing problems in computational systems biology. Recently, graph learning (GL) approaches based on graph signal processing (GSP) have been developed to infer graph topology from signals defined on graphs. However, existing GL methods are not suitable for learning signed graphs, which represent a characteristic feature of GRNs, as they account for both activating and inhibitory relationships between genes. They are also incapable of handling high proportion of zero values, which represent dropouts in single cell experiments. To this end, we propose a novel signed GL approach, *scSGL*, that learns GRNs based on the assumption of the smoothness and non-smoothness of gene expressions over activating and inhibitory edges, respectively. scSGL is then extended with kernels to take the nonlinearity of co-expressions into account and handle high proportion of dropouts. From GSP perspective, this extension corresponds to assuming smoothness/non-smoothness of graph signals in a higher dimensional space defined by the kernel. The proposed approach is formulated as a non-convex optimization problem and solved using an efficient ADMM framework.

**Results:** In our experiments on simulated and real single cell datasets, scSGL compares favorably with other single cell gene regulatory network reconstruction algorithms.

**Availability:** The scSGL code and analysis scripts are available at (*https://github.com/Single-Cell-Graph-Learning/scSGL*).

## 1 Introduction

Gene regulatory networks (GRNs) represent fundamental molecular regulatory interactions among genes that establish and maintain all required biological functions characterizing a certain physiological state of a cell in an organism (Moris *et al*., 2016). Cell type identity in an organism is determined by how active transcription factors interact with a set of cis-regulatory regions in the genome and controls the activity of genes by either activation or repression of transcription (Fiers *et al*., 2018). Usually, the relationship between these active transcription factors and their target genes characterize GRNs. Due to the inherent causality captured by these meaningful biological interactions in GRNs, genome-wide inference of these networks holds great promise in enhancing the understanding of normal cell physiology, and also in characterizing the molecular compositions of complex diseases (Saadatpour *et al*., 2014; Moignard *et al*., 2015).

GRNs can be mathematically characterized as graphs where nodes represent genes and the edges quantify the regulatory relations. GRN reconstruction attempts to infer this regulatory network from high-throughput data using statistical and computational approaches. Multiple methods encompassing varying mathematical concepts have been proposed during the last decade to infer GRNs using gene expression data from bulk population sequencing technologies, which accumulate expression profile from all cells in a tissue. These methods can be broadly classified into two groups: the first group infers a static GRN, considering steady state of gene expression, while the second group uses temporal measurements to capture the expression profile of the genes in a dynamic process. A thorough evaluation of the static and dynamic models used in bulk GRN reconstruction can be found in (Marbach *et al*., 2012; Chai *et al*., 2014).

Recent advances in RNA-sequencing technologies have enabled the measurement of gene expression in single cells. This has led to the development of several computational approaches aimed at quantifying the expression of individual cells for cell-type labelling and estimation of cellular lineages. Several algorithms have been developed to arrange cells in a projected temporal order (pseudotime trajectory) based on similarities in their transcriptional states. In parallel, several dynamic models for single cell GRN reconstruction have also been developed taking into account the estimated pseudotimes. Since single cell network reconstruction algorithms try to establish functional relationships between genes taking into account the entire population of cells, it is debatable as to whether additional knowledge regarding cell state transitions may provide any added benefits (Chen and Mar, 2018; Pratapa *et al*., 2020). In summary, direct application of bulk GRN reconstruction methods may not be adequate for single cell network inference.

The complex nature of single cell gene expression experiments brings about new computational challenges. Generally, a high percentage of genes are not expressed in single cells, giving rise to the dilemma of whether this “dropout” is due to technical noise or genuine biological variability. Several statistical methods have been designed to model this “dropout” phenomenon (Kharchenko *et al*., 2014; Finak *et al*., 2015; Risso *et al*., 2017). These zero values referred to as dropouts most likely result from biological variation and may be indicative of heterogeneity in gene expression for varying cell types (Svensson, 2020; Silverman *et al*., 2020).

A variety of algorithms for network reconstruction in scRNAseq data have been recently proposed, but most of these methods fail to outperform network estimation methods developed for bulk data or microarrays. (Pratapa *et al*., 2020; Chen and Mar, 2018). To that end, we propose a network reconstruction algorithm that learns the co-expression between genes by borrowing ideas from graph signal processing (GSP) literature. GSP provides a framework for analyzing signals defined on graphs by extending classical signal processing tools and concepts (Ortega *et al*., 2018). In many applications of GSP, the graph topology is not always available, thus it must be inferred or learned from the observed data. The major approaches to graph learning (GL) include smoothness based methods (Dong *et al*., 2019), where the graph is learned with the assumption that graph signals vary smoothly with respect to graph structure; and diffusion process based models, where the graph is learned from signals that are assumed to be graph filtered versions of random processes (Mateos *et al*., 2019). In this work, we focus on learning graphs with the smoothness assumption for the following reasons. First, smooth signals admit low-pass and sparse representations in the graph Fourier domain. Thus, the GL problem is equivalent to finding efficient information processing transforms for graph signals. Second, many graph-based machine learning tasks, such as spectral clustering, graph regularized learning etc., are developed based on the smoothness of the graph signals. Finally, smooth graph signals are observed ubiquitously in real-world applications (Mateos *et al*., 2019).

Smoothness based GL is first considered in (Dong *et al*., 2016) by modeling graph signals using factor analysis, where the transformation from factors to observed signals exploits the graph topology. By imposing a suitable prior on factors, the graph signals are modelled to have low-frequency representation in the graph Fourier domain. This analysis results in an optimization problem where a graph is learned such that variation of signals over the learned graph is minimized. Different variations of this framework with constraints on the learned topology and for handling noisy graph signals were considered in (Kalofolias, 2016; Hou *et al*., 2016; Berger *et al*., 2020; Kadambari and Chepuri, 2020; Rui *et al*., 2017). All of the previous works learn unsigned graphs with the exception of (Matz and Dittrich, 2020), where a signed graph is learned by employing signed graph Laplacian defined by Kunegis *et al*. (2010). By using signed Laplacian, Matz and Dittrich (2020) aim to learn positive edges between nodes whose signal values are similar and negative edges between nodes whose signal values have opposite signs with similar absolute values. However, this approach is not suitable when graph signals are either all positive- or negative-valued, as in the case of gene expression data.

Considering the advantages of GL approaches in learning graph topologies that are consistent with the observed signals, in this paper, we propose a novel GL algorithm for the reconstruction of GRNs. In particular, we assume gene expression data obtained from cells are graph signals residing on an unknown graph structure, which corresponds to the GRN. One important characteristic of GRNs is that they are signed graphs, where positive and negative edges correspond to activating and inhibitory regulations between genes. To this end, we propose a novel and computationally efficient signed GL approach, scSGL, that reconstructs the GRN under the assumption that graph signals admit low-frequency representation over activating edges, while admitting high-frequency representation over inhibitory edges. Biologically, this modelling implies that two genes that are connected with an activating edge have similar expressions, while two genes connected with an inhibitory edge have dissimilar expressions. Another important characteristic of scRNAseq is high proportion of dropouts. We address this issue by employing kernel functions to map graph signals to a higher dimensional space and assuming low- and high-frequency representation for these high dimensional graph signals. This mapping allows us to use kernels that are developed for zero-inflated datasets.

The remainder of the paper is organized as follows. In Section 2, we present the background on graphs and kernels. Proposed signed graph learning approach is given in Section 3. Performance of scSGL on various synthetic and real datasets are reported in Section 4. Finally, Section 5 includes discussion and concluding remarks.

## 2 Background

### 2.1 Notations

An undirected graph is denoted by *G* = (*V*,*E*) where *V* is the node set with |*V*| = *n* and *E* ⊆ *V* × *V* is the edge set with |*E*| = *m*. An edge between nodes *i* and *j* is shown by *e_ij_* and is associated with a weight *w_ij_*. If 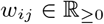, ∀*e_ij_* ∈ *E*, the graph is an unsigned graph and if 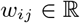, ∀*e_ij_* ∈ *E* it is a signed graph. Algebraically, *G* can be represented by an *n* × *n* symmetric adjacency matrix **W** where *W_ij_* = *W_ji_* = *w_ij_* if *e_ij_* ∈ *E* and 0, otherwise. If *G* is signed, the adjacency matrix can be decomposed into two matrices as **W** = **W**^+^ – **W**^-^ where 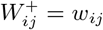 for *w_ij_* > 0 and 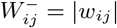 for *w_ij_* < 0. For an unsigned graph, the Laplacian matrix is defined as **L** = **D** – **W**, where **D** is the diagonal degree matrix, i.e., 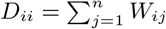. For a signed graph, we define two Laplacian matrices **L**^+^ and **L**^-^ which are constructed from **W**^+^ and **W**^-^, respectively. All-one and all-zero vectors and matrices are represented by **1** and **0**, respectively. Finally, ith row and column of a matrix **X** are represented by **X**_i_. and **X**._i_, respectively.

### 2.2 Low- and High-frequency Signals on Unsigned Graphs

A graph signal 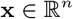 is a vector whose entries reside on the nodes of an unsigned graph *G*. Graph Fourier transform (GFT) of x is defined as the expansion of x in terms of the eigenbasis of the graph Laplacian (Shuman *et al*., 2013). This representation allows us to characterize x in terms of its graph spectral content as either low- or high-frequency, where low(high)-frequency graph signals have small (large) variation with respect to the graph (Sandryhaila and Moura, 2014).

Let **L** = **VΛV**^⊤^ be the eigendecomposition of **L** where **Λ** is the diagonal matrix of eigenvalues with Λ_*ii*_ = λ_i_ and **V**._i_ is the eigenvector corresponding to λ_*i*_. GFT of x is then 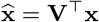 and inverse GFT is (Shuman *et al*., 2013):

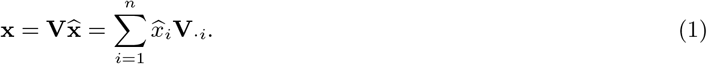

Thus, x is the linear combination of eigenvectors of **L** with the coefficients equal to the entries of 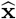. Eigenvectors of **L** corresponding to small eigenvalues have small variation over the graph. Thus, if most of the energy of 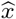 lies in 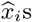 corresponding to the small eigenvalues, then x varies little over *G*. On the other hand, if most of the energy of 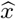 lies in 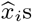 corresponding to the large eigenvalues, x has high variation over *G*. The total variation of x over *G* is then quantified as (Shuman *et al*., 2013):

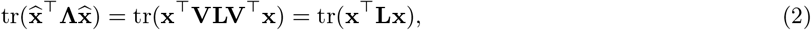

which is small for low-frequency graph signals and large for high-frequency ones.

### 2.3 Unsigned Graph Learning

An unknown unsigned graph *G* can be learned from a set of observed graph signals defined over it with the assumption that graph signals have low-frequency representation in graph spectral domain, i.e., total variation is small. Using this assumption, (Dong *et al*., 2016) proposes to learn *G* by minimizing (2) with respect to **L** given a set of graph signals 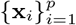 as follows:

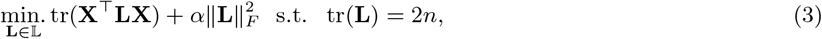

where 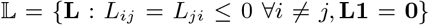 is the set of Laplacian matrices and 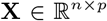 is the data matrix whose columns are graph signals. ||**L**||_*F*_ is the Frobenius norm introduced to control the sparsity of the learned graph such that larger *α* values result in denser graphs. Finally, the constraint tr(**L**) = 2*n* ensures that trivial solutions are avoided.

### 2.4 Kernels

Traditional machine learning and signal processing applications are mostly developed based on linear modelling due to their simplicity. However, real world problems require nonlinear estimation that can detect more complex patterns in the data. For this purpose, kernels are introduced to capture the nonlinearity by mapping signals to a high-dimensional space (Hofmann *et al*., 2008). Kernels correspond to dot products in a higher dimensional feature space and overcome explicit construction of the feature space; thus providing simplicity of linear methods in nonlinear estimation. Given data from input space 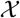, and a mapping function 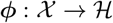 where 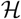 is an Hilbert space, a kernel function can be expressed as an inner product in the corresponding feature space, i.e., 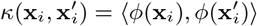, where 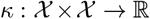 is a finitely positive semi-definite kernel function (Shawe-Taylor *et al*., 2004). An explicit representation of the feature map *ϕ* is not necessary and the dimension of mapped feature vectors could be high and even infinite. By using different kernels, learning algorithm can be augmented to exploit various (nonlinear) associations between input data. For example, the first term in (3) can be rewritten as tr(**XX**⊤**L**) = ∑_*i,j*_〈**X**_*i*_., **X**_*j*_.〉*L_ij_*, where the dot product between the rows of **X** can be replaced by *κ*(**X**_*i*_., **X**_*j*_.). In the next section, this observation is used to develop a graph learning framework that is able to capture nonlinear relations between graph signals.

## 3 Methods

### 3.1 Signed Graph Learning

In (3), an unsigned graph is learned with the assumption that the observed graph signals have low-frequency representation in graph spectral domain. In order to learn a signed graph *G*, one needs to make some additional assumptions about the graph signals **X**. In this work, we make the following assumptions:

1. Signal values on nodes connected by positive edge values are similar to each other, i.e., variation over positive edges is small.
2. Signal values on nodes connected by negative edge values are dissimilar to each other, i.e., variation over negative edges is large.

From GSP perspective, these assumptions correspond to graph signals being low- and high-frequency over positive and negative edges, respectively. Let *G*^+^ be the graph corresponding to the positive edges of *G* and let *G*^-^ be the graph corresponding to the negative edges of *G* with the edge weights equal to the absolute value of the original edge values. Assumption 1 implies that the graph signals have low-frequency representation in the graph Fourier domain of *G*^+^. On the other hand, assumption 2 implies that the graph signals have high-frequency representation in graph Fourier domain of *G*^-^. We use (2) to quantify how well the graph signals fit these assumptions. Thus, to learn an unknown signed graph, we minimize tr(**X**^⊤^**L**^+^**X**) with respect to **L**^+^ while maximizing tr(**X**^⊤^**L**^-^**X**) with respect to **L**^-^:

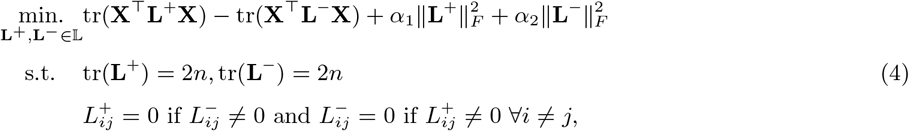

where Frobenius norms and the first two constraints are similar to (3) and the last constraint ensures that *L*^+^ and *L*^-^ are not non-zero for the same indices.

### 3.2 Kernelized Signed Graph Learning

As mentioned in Section 2.4, kernels are used in learning algorithms to exploit various (nonlinear) relations between input data. This is especially crucial in GRN inference as shown in (Skinnider *et al*., 2019), where 17 different association measures between gene expressions are compared in terms of their performance in GRN inference and various other tasks on single-cell transcriptomic datasets. In this paper, we consider three kernels: correlation coefficient, *r*, measure of proportionality, *ρ* (Quinn *et al*., 2017) and a modification of Kendall’s tau (*τ_zi_*) for zero inflated non-negative continuous data (Pimentel *et al*., 2015). These kernels are selected because *r* is a commonly used measure for network inference, *ρ* performs the best in (Skinnider *et al*., 2019) and *τ_zi_* can handle high ratio of dropouts in scRNAseq. In its current form, (3) cannot be used directly for different associations. Thus, the optimization problem in (4) is extended using kernels. The first term in (4) can be written as 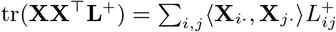 and the second term can be written similarly. By replacing dot products with a given kernel function, i.e., *κ*(**X**_i_., **X**_j_.), the problem in (4) can be extended to incorporate the different associations as:

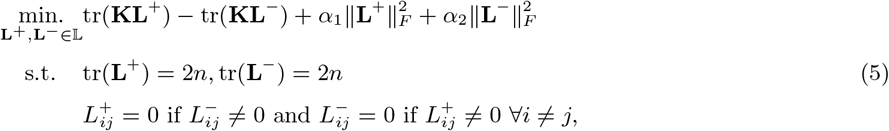

where 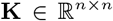 is the kernel matrix with *K_ij_* = *κ*(***X**_i_*., **X**_*j*_.). From GSP perspective, this modification implies that graph signals on each node, i.e., **X**_*i*_., are first mapped to a (higher dimensional) Hilbert space and the signed graph is learned in this new space. The graph signals in this space, i.e., [*ϕ*(**X**_1_.)_j,…,_ *ϕ*(**X**_*n*_.)_j_]^⊤^ where *ϕ*(**X**_*i*_.)_*j*_ is the *j*th entry of the vector *ϕ*(**X**_*i*_.), are assumed to have low- and high-frequency representation with respect to *G*^+^ and *G*^-^, respectively. Extending signed graph learning problem in (4) using kernels brings flexibility and any association metric in (Skinnider *et al*., 2019) can be implemented in this framework if it is a positive semi-definite kernel. The optimization procedure for (5) is given in the Supplementary Material.

### 3.3 Hyperparameter Selection

The optimization problem in (5) requires the selection of two regularization parameters *α*_1_ and *α*_2_, which determine the density of the learnt graph, i.e., large values of *α*_1_ (*α*_2_) result in denser **L**^+^ (**L**^-^). Their values can be set to obtain a graph with desired positive and negative edge densities. In supplementary material, a resampling approach (Efron and Tibshirani, 1986) is provided to determine positive and negative edge densities empirically. This approach is followed for all datasets analyzed in the next section.

## 4 Results

In this section, performance of scSGL is evaluated and compared to state-of-the-art GRN inference methods on various simulated and experimental scRNAseq datasets. We selected GENIE3 (Huynh-Thu *et al*., 2010), GRNBOOST2 (Moerman *et al*., 2019), PIDC (Chan *et al*., 2017) and PPCOR (Kim, 2015) for comparison as they are the top performing methods in (Pratapa *et al*., 2020). GENIE3, GRNBOOST2 and PPCOR were originally developed for bulk analysis, while PIDC is developed for single cell gene expression data. Among these methods, GENIE3 and GRNBOOST2 return fully connected directed networks, while the remaining two infer undirected networks. Finally, only PPCOR algorithm returns signed graphs. Given the inherent sparsity of gene networks, we used the area under the precision-recall curves (AUPRC) as the primary evaluation metric. Results using area under the receiver operating characteristic curves (AUROC) and early precision ratio (EPR) are reported in Supplementary Material.

### 4.1 Synthetic Datasets

#### Curated Datasets From BEELINE

The first simulation datasets we consider are curated from ‘‘published Boolean models of GRNs” (Pratapa *et al*., 2020). These datasets were generated using the recently proposed single cell GRN simulator BoolODE (Pratapa *et al*., 2020). BoolODE converts boolean functions specifying a GRN directly to ODE equations using GeneNetWeaver (Schaffter *et al*., 2011; Marbach *et al*., 2009), a widely used method to simulate bulk transcriptomic data from GRNs. These datasets are generated from four literature-curated Boolean models: mammalian cortical area development (mCAD), ventral spinal cord (VSC) development, hematopoietic stem cell (HSC) differentiation and gonadal sex determination (GSD). These models represent different types of graph structures, with varying numbers of positive and negative edges; thus serving as good examples for illustrating the robustness of the proposed method in modelling signed graph topologies. BoolODE is used to create ten random simulations of the synthetic gene expression datasets with 2,000 cells for each model. For each dataset, one version with a dropout rate of 50% and another with a rate of 70% are also considered to evaluate the performance of the methods under missing values.

AUPRC values are calculated separately for activating and inhibitory edges and their average over realizations are reported in Figure 1. For most of the datasets, scSGL performs better than other benchmarking methods in inferring both activating and inhibitory edges. Although there is a difference between the performances of different kernels, scSGL generally performs better than state-of-the-art methods irrespective of the selected kernel. Comparing the performances of different kernels, it is observed that *τ_zi_* results in higher AUPRC values in GSD, HSC and VSC while *ρ* performs better in mCAD datasets. Increasing the dropout ratio causes a drop in the performance of all methods for inferring the activating edges but not for learning the inhibitory edges. Overall, the best performing kernel is *τ_zi_*, which might be because of its robustness to increasing dropout ratio compared to other kernels.

**Figure 1:**
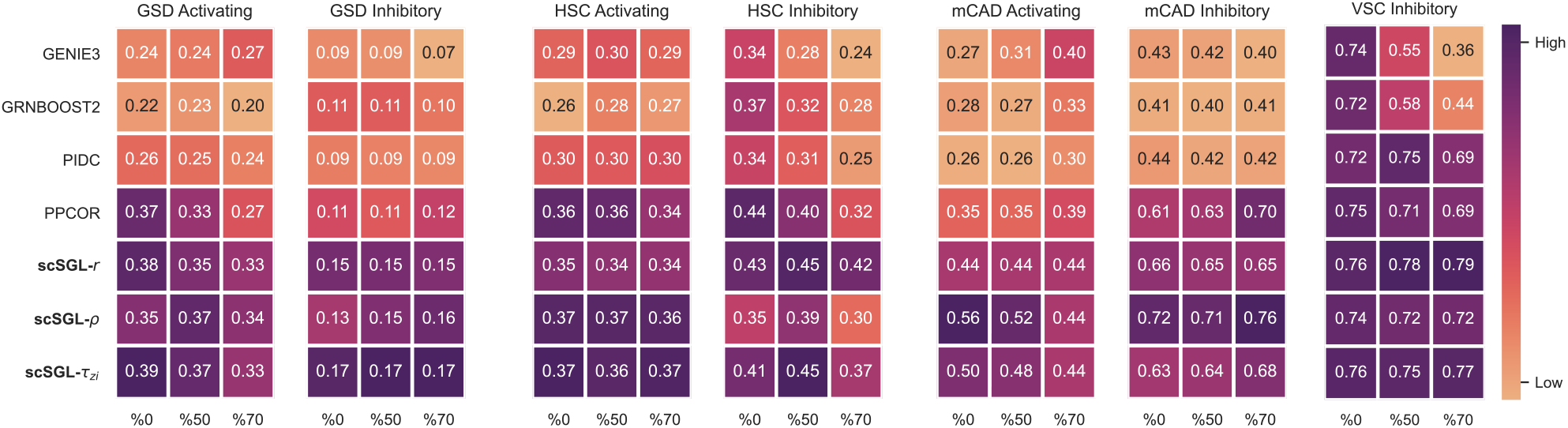
Performance of scSGL and state-of-the-art methods on curated datasets as measured by AUPRC for activating and inhibitory edges. x-axis indicates dropout ratio in the dataset.

#### Parameter sensitivity analysis

To mimic the zero inflated and overly dispersed nature of most scRNAseq datasets, we simulated gene expression data from a multivariate zero-inflated negative binomial (ZINB) distribution for our second simulation. These datasets were then used to conduct parameter sensitivity analysis for the proposed methods. Given a known graph structure, synthetic datasets are generated from a ZINB distribution by adapting an algorithm developed by (Yahav and Shmueli, 2012). The three parameters of the ZINB distribution; λ, *κ* and *ω*, which control its mean, dispersion and degree of zero-inflation, respectively were determined from a real scRNAseq dataset to make the simulations mirror the properties of real datasets (See Supplementary Material).

The ZINB simulator is then used to generate expression data from three different network structures: random networks, networks with a given community structure and networks with hubs. Random networks are generated using Erdős-Rényi model with desired edge density. Since Erdos-Rényi model is not realistic due to its binomial degree distribution, we also consider networks with hubs. These networks are generated using a Barabési-Albert model whose degree distribution follows a power-law function. Finally, networks with community structure, also known as modular networks, are generated using a disjoint union of random graphs. The accuracy of the scSGL inferred graphs were then evaluated for all three graph structures. To investigate the robustness of scSGL, we simulated datasets from the aforementioned network topologies by varying the following parameters: (*i*) number of genes (10, 50, 100 and 250), (*ii*) number of cells (100, 300, 500 and 1000) and (*iii*) dropout probabilities (0.26-0.36). To account for the inherent randomness of the simulations, 10 independent data replicates were generated for every parameter combination and the mean AUPRC scores obtained by averaging over the replicates are reported in Figure 2.

**Figure 2:**
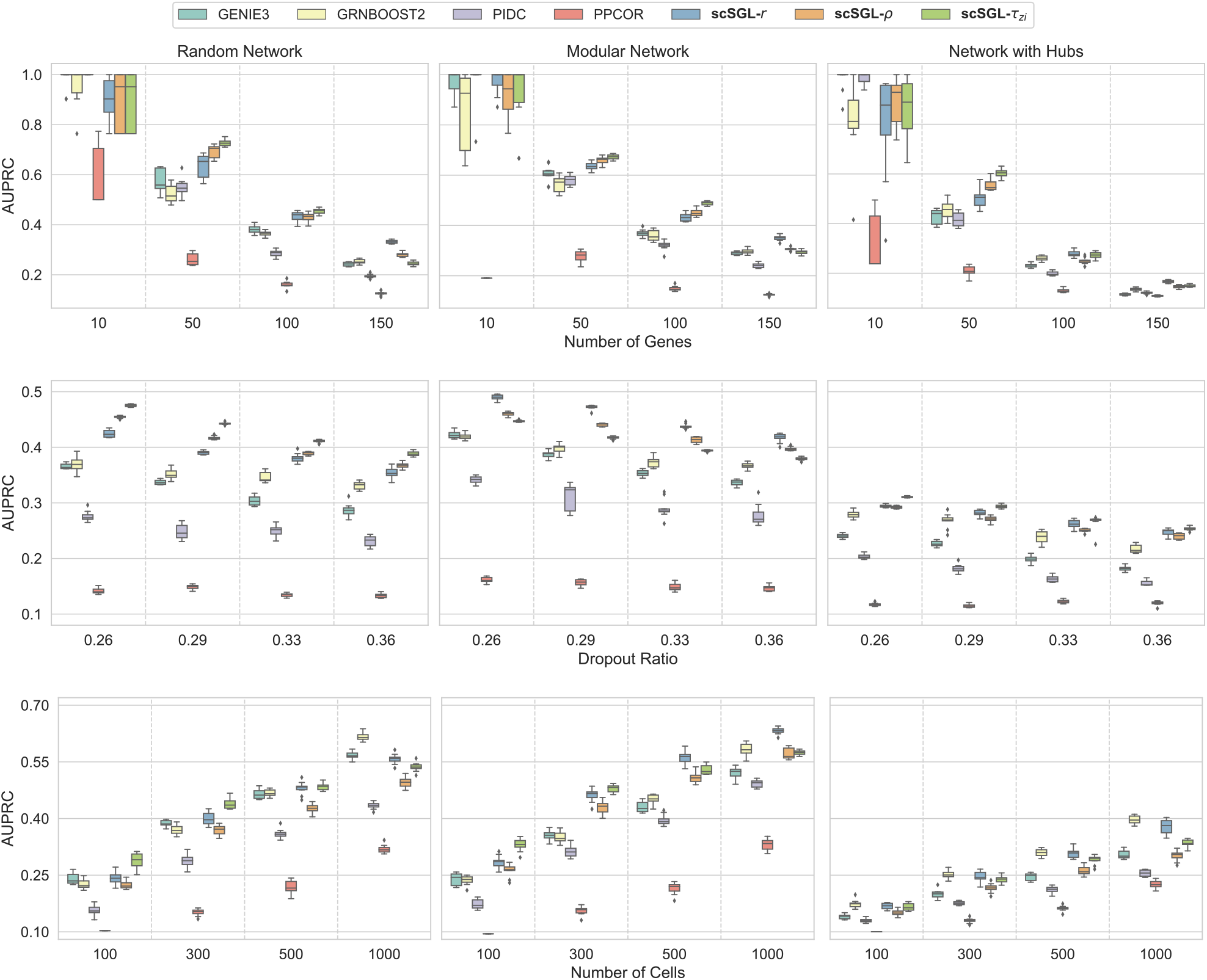
Performance of various methods for synthetic datasets with varying number of genes (top row), dropout ratio (middle row) and number of cells (bottom row).

Recent investigations of scRNAseq datasets have revealed that dropout rates are primarily driven by a combination of technical and biological factors (Fiers *et al*., 2018). Consequently, while mean gene expression and proportion of zeros are linked, this may vary based on cell type, sex, and other biological and technical factors. While investigating the impact of dropout rates on network estimation accuracy, we found a steady decline in AUPRC scores for all methods with an increase in the number of zeroes. scSGL irrespective of the kernel choice maintained the highest AUPRC scores across all network topologies. Gene expression in scRNAseq datasets can be intepreted as relative measures of abundance owing to the datasets being a combination of gene expression derived from several celltypes. This could be the reason why proportionality measures perform well (Skinnider *et al*., 2019). The strong performance of *τ_zi_* can be explained on the basis that it explicitly models the dropouts present in scRNAseq datasets. Despite the poor performance of regularized correlation networks (PPCOR), we see a strong performance of scSGL when using the correlation kernel. This proves that gene-gene relationships are in fact non-linear in nature. This belief is also strengthened by the above average performance of tree-based machine learning algorithms like GENIE3 and GRNBOOST2. It is to be noted that PIDC, the only other method capable of modelling excess zeroes, while accounting for non-linear relationships fails to achieve a top-ranking AUPRC score.

Next, we evaluated the impact of cell sizes on network reconstruction. Figure 2 demonstrates a clear rise in AUPRC scores when the number of cells are increased. PIDC, the only other single cell network estimation technique, achieves a below average performance at the lowest sample size of 100. This could be due to the fact that PIDC requires large sample sizes for accurate estimation of pairwise joint probability distributions for calculating mutual information. In general, PPCOR has the worst performance among all methods. It should also be noted that the performance of GRNBOOST2 was equivalent to scSGL for all the network topologies when the sample size was 10 times the number of genes. These results indicate the importance of sample size in accurate network estimation for all of the methods and network topologies is considered.

Finally, the performance of each of the methods was evalulated by varying the number of genes. All methods had high AUPRC across network topologies when the number of genes was small. While the AUPRC scores of all the methods declined with an increase in the number of genes, scSGL performed significantly better than most of the benchmarking methods. This dip in performance could be attributed to the fact that all methods learn very dense networks. With an increase in the number of nodes, there is an increasing number of false edges detected by every algorithm. The performance of scSGL could further be improved with a more biologically informed framework for hyperparameter selection.

### 4.2 Real Datasets

For real datasets, we consider scRNAseq expressions of human embryonic stem cells (hESC) and mouse embryonic stem cells (hESC) which include 758 and 451 cells, respectively. We inferred GRNs between 500 highly varying genes along with highly varying TFs (Pratapa *et al*., 2020). Inferred GRNs are compared to three different databases of gene regulations: STRING (Szklarczyk *et al*., 2021), cell-type specific (Consortium *et al*., 2012) and nonspecific (Liu *et al*., 2015; Garcia-Alonso *et al*., 2019; Han *et al*., 2018). AUPRC values are calculated for each method and the ratios to AUPRC of the random estimator are reported in Figure 3. Although all methods have performance values close to random estimator, scSGL performs better than state-of-the-art methods in hESC and has comparative performance in mESC dataset.

**Figure 3:**
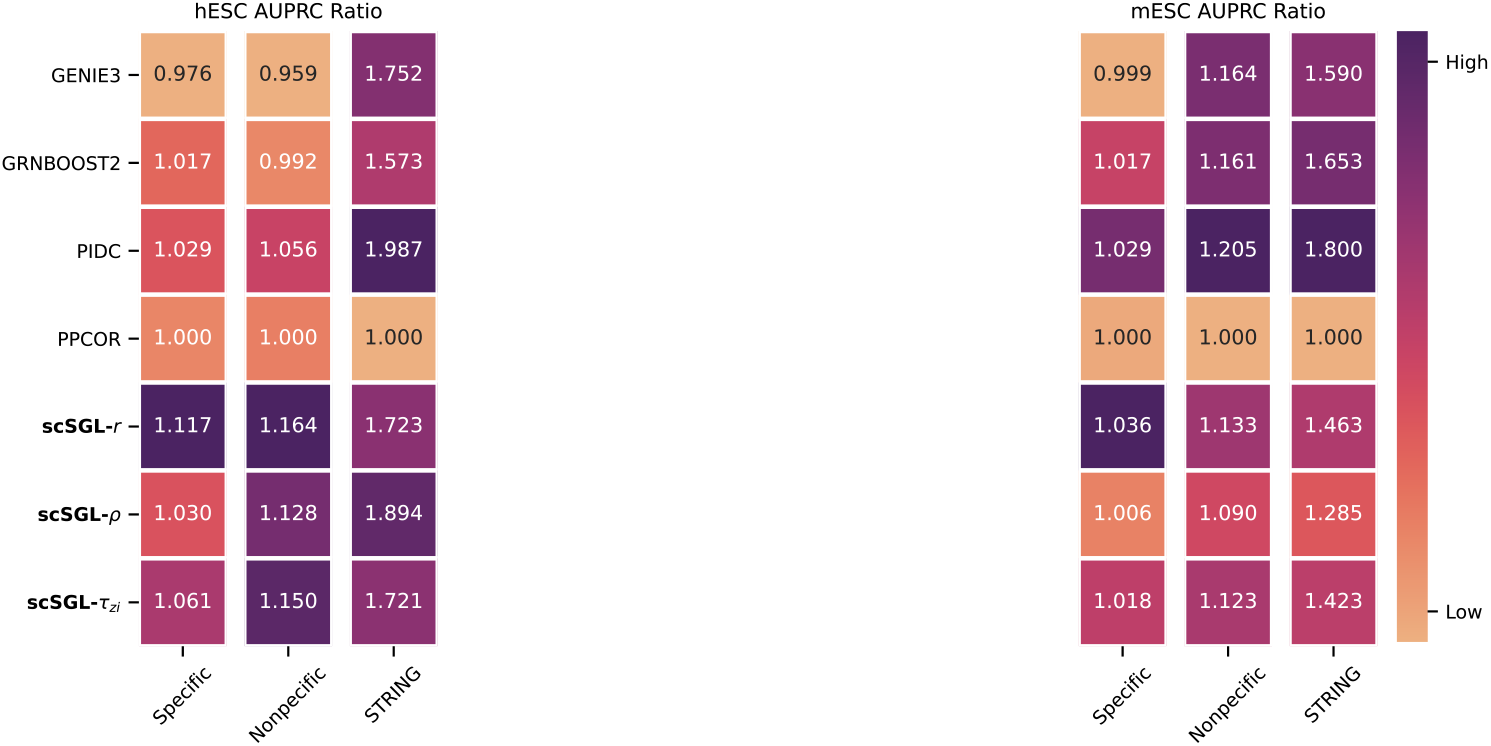
Performance of methods for two real-world scRNAseq datasets. Inferred graphs are compared to three different gene regulatory databases.

To add biological meaning to the estimated networks we compared them to the reference networks in the STRING database. The STRING database is a compendium of protein-protein interactions created by gathering information from varying sources like experimental studies, text mining etc. The edges in the STRING network are classified as high confidence (minimum score of 0.700), medium confidence (minimum score of 0.400) and and low confidence (minimum score of 0.150). In hESC dataset, scSGI.-*ρ* identified the maximum number of high confidence associations present in the STRING reference network. scSGL-*ρ*, scSGL-*r* and scSGL-*τ_zi_* each identified 60, 56 and 24 high confidence STRING interactions, respectively, with an edge confidence greater than 0.5. The interactions identified by scSGL-*r* form a network of 56 unique genes including genes Nanog, Sox2, Sox4, Pou5f1, Ctnnb1, Gata2, Gata3 and many others. Lineage-specific marker genes, Cdk6, Col5a1, Vim, and Itg5, which are known to have regulatory roles in cell differentiation were also detected by scSGL-*r* but with edge confidence less than 0.5 (0.1-0.3) (Brafman *et al*., 2013; Chu *et al*., 2016). scSGL-*ρ* and scSGL-*r* identified 20 common genes including Sox2, Sox4, Gata6, Ctnnb1 and Bmp4. scSGL-*τ_zi_* identified the least number of genes but successfully retrieved lineage markers Nanog, Sox2, Sox4, Pou5f1,Ctnnb1, Gata2, Gata3. All three kernel methods identified genes Sox4, Ctnnb1, Bmp4 and Gata6. According to the STRING database, the 56 genes identified by scSGL-*r* are associated with 839 significantly enriched biological process gene ontology (GO) terms that include cell differentiation, chromosome separation, specification of animal organ position, mitotic nuclear division and organ formation. Genes identified by scSGI.-*ρ* and scSGL-*τ_zi_* had similar functional enrichments for biological processes.

In mESC dataset, scSGL-*ρ*, scSGL-*r* and scSGL-*τ_zi_* each identified 67, 103 and 55 high confidence STRING interactions, respectively, with an edge confidence greater than 0.5. The three estimated networks capture interactions regulated by known transcription factors Sox2, Nanog, Klf4, Myc and Sall4 (Zhou *et al*., 2007). scSGL-*r* identified known relationships between Sox2 and Nanog; Esrrb with Sox2 and Rybp among many others. scSGI.-*ρ* identified known relationships between Esrrb and Etv5 and indirect interactions between Sall4 and Rybp regulated by TF Oct4. scSGL-*τ_zi_* identified most of the important relationships identified by scSGL-*r* along with additional relationships between Sox2, Nanog, and Rif1. According to the STRING database, the 103 genes identified by scSGL-r are associated with 908 significantly enriched biological process GO terms that include cell fate determination, specification and commitment, mitotic DNA replication and regulation of nodal signalling pathway. Similar to hESC analysis, scSGL, irrespective of the chosen kernel, identified genes with similar functional enrichments for biological processes.

Finally, to analyze the relation between edges identified by scSGL and benchmarking methods, the intersection between the top 1000 edges is reported as an UpSet plot (Lex *et al*., 2014) in Figure 4. In both datasets, PPCOR does not have any intersection with other methods probably because of its poor performance reported in Figure 3. The remaining 6 methods have an intersection set with cardinality around 40 edges. The same number of common edges is found in the intersection of PIDC, GENIE3, GRNBOOST2, scSGL-*τ_zi_*, scSGL-*r* and in the intersection of PIDC, GENIE3, scSGL-*τ_zi_*, scSGL-*r*, scSGL-*ρ*. These observations hold for both datasets, indicating the reproducibility of the proposed approach across different datasets. Edges identified by *τ_zi_* and *r* have more intersecting edges with benchmarking methods and with each other than those identified by *ρ*, which indicates that the benchmarking methods have more common edges with correlation based association metrics than with proportionality measures. scSGL methods have more common edges with PIDC than with GENIE3 and GRNBOOST2, which may be due to the fact that PIDC learns co-expression GRN similar to scSGL, while GENIE3 and GRNBOOTS2 learn directed interactions between genes.

**Figure 4:**
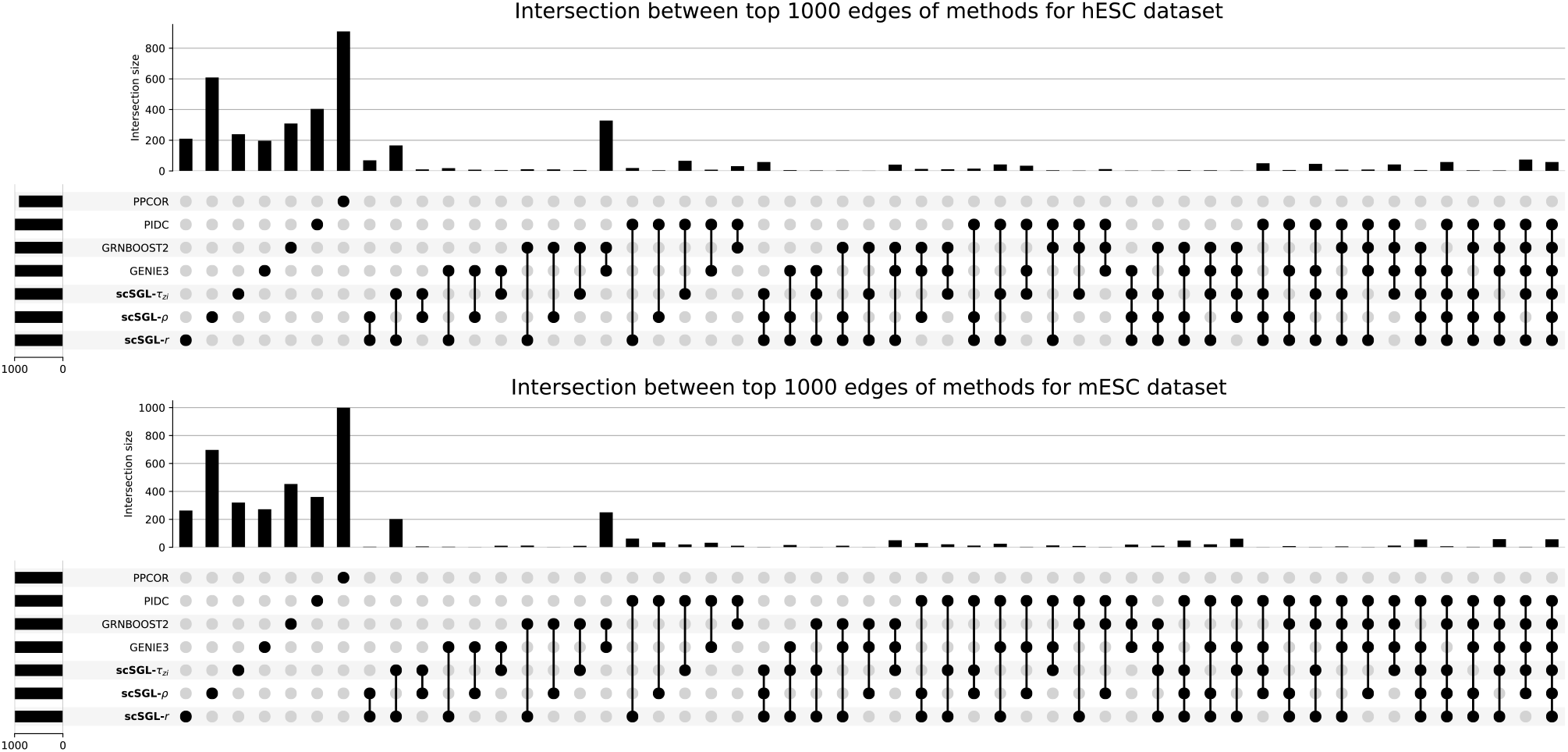
UpSet plot that shows intersection between the top 1000 edges by scSGL with 3 kernels and benchmarking methods in hESC and mESC datasets.

## 5 Discussion and Conclusion

In this paper, we have introduced a novel network inference algorithm based on GSP. Our proposed algorithm scSGL identifies functional relationships between genes by learning the signed adjacency matrix from the gene expression data under the assumption that graph signals are similar over positive edges and dissimilar over negative edges. This novel technique also takes into account the nonlinearity of the gene interactions by employing kernel mappings. We applied scSGL to four curated datasets derived from ‘‘published Boolean models of GRN” and two real experimental scRNAseq datasets during differentiation. To conduct an in-depth analysis of gene co-expression network reconstruction from scRNAseq datasets, we generated simulations from zero inflated negative binomial distributions. These simulations, generated using different parameter combinations, were used to investigate the robustness of our proposed methods to changing cell sizes, gene numbers and dropout rates.

For the curated datasets, scSGL consistently obtained higher AUROC and AUPRC scores in comparison to the benchmarking methods, despite each dataset having a different number of stable cell states. Parameter sensitivity analysis reflected the superior performance of scSGL in estimating networks under varying network topologies. The performance remained consistent even when the gene numbers increased, the dropout rates were high and the sample sizes were low. This indicated the robustness of scSGL in modelling networks under varying characteristics of scRNA-seq datasets.

The networks estimated from real data using scSGL identified important functional relationships between target genes and transcription factors and exhibited enrichment for appropriate functional processes. We also demonstrated that scSGL attained performance comparable to state-of-the-art-methods in real data experiments, with the performance of all the GRN reconstruction methods methods being close to random. Accuracy evaluation of the predicted networks for the real datasets were done using cell-type specific, non-specific and functional networks described in (Pratapa *et al*., 2020). However, most of the information in these ground truth datasets have been accumulated based on tissue level data and hence it’s not completely appropriate to calculate precision and recall rates from these databases.

Although scRNAseq techniques provide significant advantages over bulk data such as increased sample size with higher depth coverage and and presence of highly distinct cell clusters, it also comes laced with multiple sources of technical and biological noise. Moreover, the inability to differentiate between technical and biological noise, and the absence of adequate noise modelling techniques further exacerbate the problem (Grün *et al*., 2014; Stegle *et al*., 2015). scSGL aims to capture the node similarities and dissimilarities based on distances between graph signals. These graph signals exhibit smoothness, which implies that within a given node cluster, genes tend to be homogeneous, while varying across clusters. This leads to densely connected graphs where the heterogeneity induced by distinct cell sub-populations can be simultaneously curbed. Using single cell data with cell cluster labels, easily obtained from single cell clustering algorithms (Petegrosso *et al*., 2020), in conjunction with scSGL can aid in identifying functional modules that are associated with a cell type (Wang *et al*., 2020). Integrating pseudotemporal ordering with scSGL can further help in identifying the functional modules associated with differential pathways (Xue *et al*., 2013).

Despite the availability of a large number of computational methods, accurate GRN reconstruction still remains an open problem. Most reconstruction methods are based on the assumption that presence of an edge implies regulatory relationships. They also have the tendency to establish links between genes regulated by the same regulator. These issues can generate a lot of false positives and therefore additional sources of data such as ChIP-seq measurements that help in identifying direct interactions between TFs and target genes, can provide a way to filter out the spurious interactions (Aibar *et al*., 2017). Finally, gene regulation has multiple layers beyond direct TF-target interaction, but functional relationships can only be established if these relationships induce persistent changes in transcriptional state. As single cell data sources over multiple modalities continue to become available, it will be interesting to see how integration of these data types aids GRN reconstruction using scSGL. (Stuart *et al*., 2019).

## 6 scSGL - Supplementary Material

### 1 Optimization Algorithm for Signed Graph Learning

In this section, we present an ADMM based algorithm to solve the optimization problem for signed graph learning. For convenience, we include the optimization problem below:

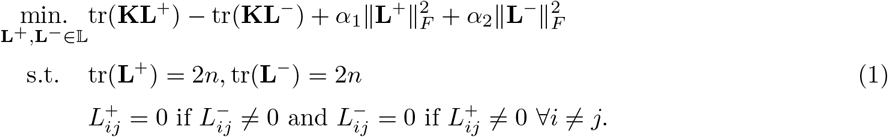

This problem is non-convex due to the last two constraints, which are called complementarity constraints (Scheel and Scholtes, 2000). In (Wang *et al*., 2019), it is shown that alternating direction method of multipliers (ADMM) converges for problems with complementarity constraints under some assumptions. First, we rewrite the problem in vector form. Let upper(·) be an operator that takes an *n* × *n* matrix and returns a *n*(*n* – 1)/2-dimensional vector that corresponds to the upper triangular part of the input matrix. Define diag(x) as an operator which returns a diagonal matrix with the diagonal elements equal to the input vector x. Similarly, diag(**X**) returns the diagonal of the input matrix **X** as a vector. The matrix 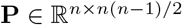 is defined such that **P**upper(**A**) = **A1** where **A** is a symmetric matrix whose diagonal entries are equal to zero. Let k = upper(**K**), **d** = diag(**K**), *ℓ*^+^ = upper(**L**^+^), *ℓ*^-^ = upper(**L**^-^). Thus, (1) can be rewritten as:

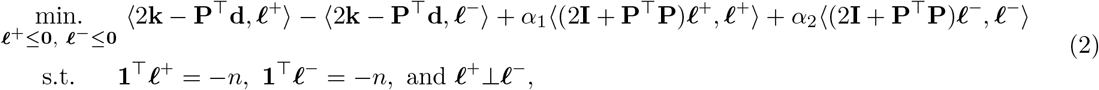

where the first two terms correspond to trace terms in (1), the last two terms correspond to Frobenius terms of (1) and first two constraints are the same as the first two constraints of (1). The last constraint with *ℓ*^+^ ≤ **0** and *ℓ*^-^ ≤ **0** correspond to the complementarity constraints. By introducing two slack variables v = *ℓ*^+^ and w = *ℓ*^-^, the problem is written in standard ADMM form:

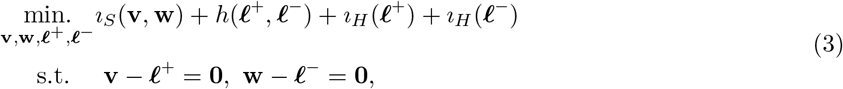

where *i_s_*(·) is the indicator function for the complementarity set *S* = {(**v**, **w**): **v** ≤ 0, **w** ≤ 0, **v**▥**w**}, *h*(*ℓ*^+^, *ℓ*^-^) is the objective function in (2), and *i_H_*() is the indicator function for the hyperplane *H* = {*ℓ*: **1**^⊤^*ℓ* = – *n*}. The augmented Lagrangian of (3) is:

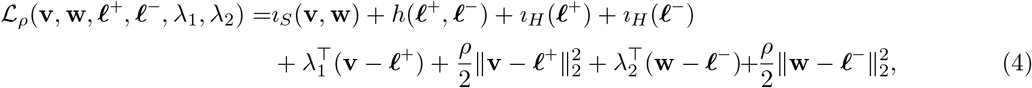

where λ_1_ and λ_2_ are Lagrange multipliers and *ρ* > 0 is the Augmented Lagrangian parameter.

#### (v, w)-step

The (**v**, **w**)-step of ADMM can be found as the projection onto the complementarity set S:

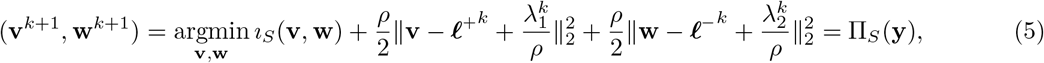

where 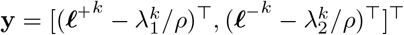 and Π_*S*_(·) is the projection operator on the set *S*.

#### (*ℓ*^+^, *ℓ*;^-^)-step

Using the fact that optimization can be performed separately for *ℓ*^+^ and *ℓ*^-^, *ℓ*^+^-step can be written as:

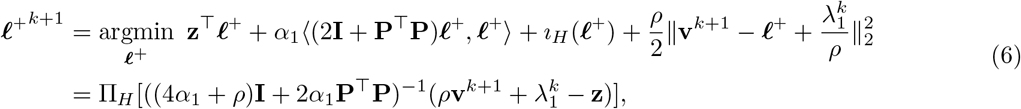

where z = 2k – P^⊤^d and Π_H_(·) is the projection operator on the hyperplane *H*. Similarly, *ℓ*^-^-step can be written as:

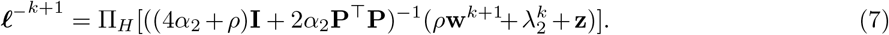

#### Lagrange multipliers udpate

The updates of Lagrange multipliers are:

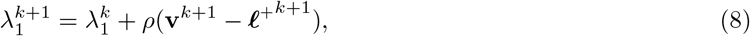

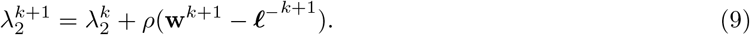

### 2 Hyperparameter selection

The optimization problem in (1) requires the selection of two regularization parameters *α*_1_ and *α*_2_, which determine the density of the learnt graph, i.e. large values of *α*_1_ (*α*_2_) result in denser **L**^+^ (**L**^-^). Thus, *α*_1_ and *α*_2_ can be set to the value that returns a graph with desired density, which we propose to determine empirically using the following procedure:

1. Given a matrix 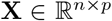 whose columns are graph signals, we randomly shuffle each column of the matrix *k* times creating *k* surrogate data matrices.
2. Association between rows of the surrogate data matrices are calculated by the kernel employed in (1).
3. Thresholds λ_1_ and λ_2_ are selected as the *p*th and (100 – *p*)th percentiles of the values in the association matrix calculated in Step 2.
4. Steps (1-3) are repeated *k* times to construct the empirical distribution of the thresholds λ_1_ and λ_2_.
5. Finally, 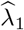 and 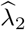 are selected to be the medians of the empirical distributions constructed in Step 4.
6. The association matrix for the original data **X** is constructed.
7. The number of entries in the association matrix that are smaller than 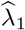 are determined and normalized by the total number of entries in the association matrix to obtain the density of **L**^-^. Similarly, number of entries in the association matrix greater than 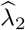 is used to determine the density of **L**^+^.
8. Values of *α*_1_ and *α*_2_ are then selected to learn graphs with the estimated graph densities found in Step 7. Since the density of the positive (negative) graph increases monotonically with the value of *α*_1_ (*α*_2_), bisection search is used to determine the values of *α*_1_ and *α*_2_ that give the desired densities.

#### Remark

For all the datasets analyzed in the main text, we learned the densities of positive and negative parts by setting *p* = 5.

### 3 Generation of simulated datasets from zero-inflated negative binomial distribution

In this section, we outline the algorithm that was used to generate the simulated datasets for parameter sensitivity analysis.

1. For each simulation setting, we first generated a binary association graph *G*_*j*1*j*2_, ∀*j*_1_,*j*_2_ ∈ {1,…,*p*} using either of the three graph topologies random, hub and cluster.
2. A binary indicator *I*_*j*1*j*2_ was next sampled for each entry of the association graph with *I*_*j*_1_*j*2_ ~ *Bernoulli*(0.5).
3. Given the binary association graph a weight matrix was generated as:

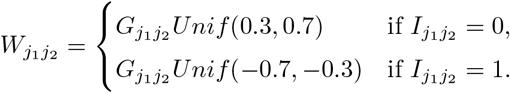
4. Random samples of *n* multivariate Gaussian random variables were then generated, with known weight matrix *W*_*j*1*j*2_. The random sample was denoted as (*X*_1_,…,*X_p_*), where each variable (gene vector) *X_j_* = (*X*_*j*1_,…, *X_jn_*)^*T*^ consisted of n realizations.
5. To mimic the dropout phenomenon present in real scRNAseq datasets, we next introduced additional zeros to the gene expression matrix. Following (Pierson and Yau, 2015), the dropout probability for each row (gene vector) in the gene expression matrix **X** was calculated as: 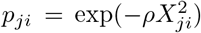, where *ρ* represents the exponential decay parameter that controls the dependence between the dropout probability and gene expression.
6. A binary indicator was next sampled for each entry: *η_ji_* ~ *Bernoulli*(*p_ji_*), with *p_ji_* = 1 indicating that the corresponding entry of *X_ji_* would be replaced by 0. The dropout probability for each gene vector was calculated as 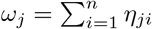.
7. Using a modification of the NORTA (Normal to Anything) method (Yahav and Shmueli, 2012) we generated samples from a multivariate zero inflated negative binomial distribution based on the multivariate normal samples generated in Step 4 using mean, dispersion and zero-inflation parameters λ, *k* and *ω_j_s*.
8. To mirror real scRNA-seq gene expression data behaviour, the gene expression mean λ and standard deviation *k* were estimated from a real scRNA-seq dataset, Peripheral Blood Mononuclear Cells (PBMC) freely available from 10X Genomics.

### 4 Performance Metrics

#### AUPRC and AUROC

Area under precision and recall curve and area under the receiver operating characteristic are calculated by comparing inferred graphs to ground truth gene regulations. During this calculation, signs of the learned edges are ignored as the AUPRC and AUROC are performance metrics restricted to binary classification. In particular, we first take the absolute value of edge weights and then compare them to ground truth edges. Thus, these metrics indicate how well methods detect edges without considering the signs of the inferred edges. Ground truth networks are considered as undirected and self-loops are ignored. Following (Pratapa *et al*., 2020), we also defined *AUPRC ratio* and *AUROC ratio* as the ratio of AUPRC (AUROC) value of the methods to AUPRC (AUROC) of the random estimator.

#### AUPRC Activating/Inhibitory

One of our goals is to learn whether the edges are activating or inhibitory. AUPRC as defined above cannot evaluate the sign information. Thus, for curated datasets, whose ground truth gene regulations include signed edge information, we calculate AUPRC for activating and inhibitory edges seperately. In particular, for methods that learn signed graphs we compare the learned positive edges to activating edges in ground truth and learned negative edges to inhibitory edges in the ground truth. For methods that do not learn signed edges, we compare all inferred edges to the ground truth activating and inhibitory edges separately to calculate their AUPRC activating and inhibitory values.

#### EPR

Early precision ratio is the fraction of true positives in the top-*k* edges in the inferred graphs where *k* is the number of edges in the ground truth network (Pratapa *et al*., 2020). For methods that return signed edges, we found top-k edges after taking absolute value of the edge weights, thus this metric is used for edge detection performance rather than the detection of edge signs. Ground truth networks are considered to be directed graphs and self-loops are ignored. Finally, *EPR ratio* is defined as the ratio of the EPR values of the methods to the EPR value of the random estimator.

### 5 AUROC and EPR Results

In the main text, we consider AUPRC based metrics defined above as the main performance metrics for comparison due to inherent sparsity of GRNs. In this section, AUROC and EPR values for synthetic data used in parameter sensitivity analysis are reported in Figures 1 and 2, respectively. AUROC and EPR ratios for real datasets are also reported in Figure 3.

---

**Figure 1:**
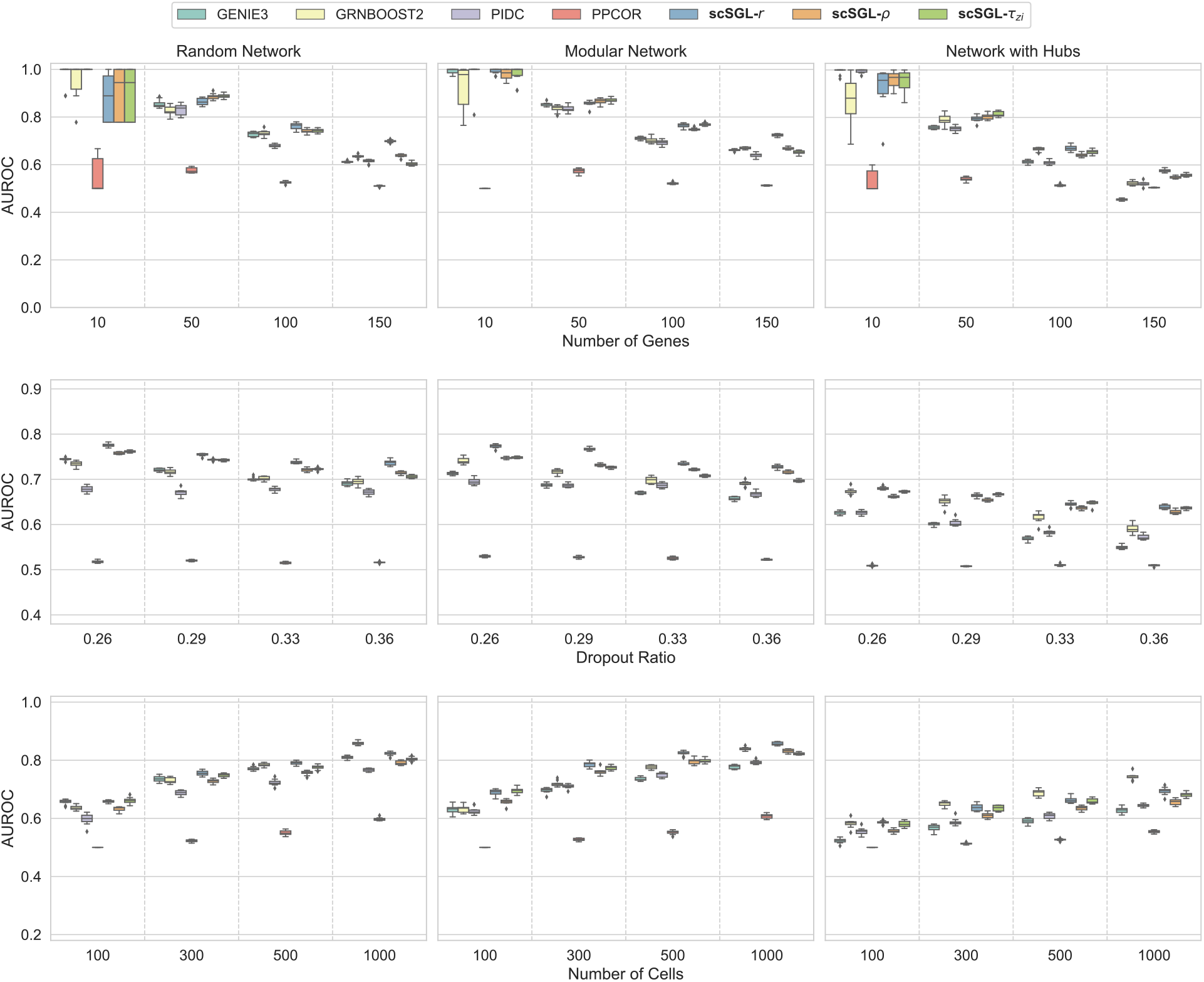
AUROC values of various methods for synthetic datasets with three different topologies (random, modular and hub) and varying number of genes (top row), dropout ratio (middle row) and number of cells (bottom row).

**Figure 2:**
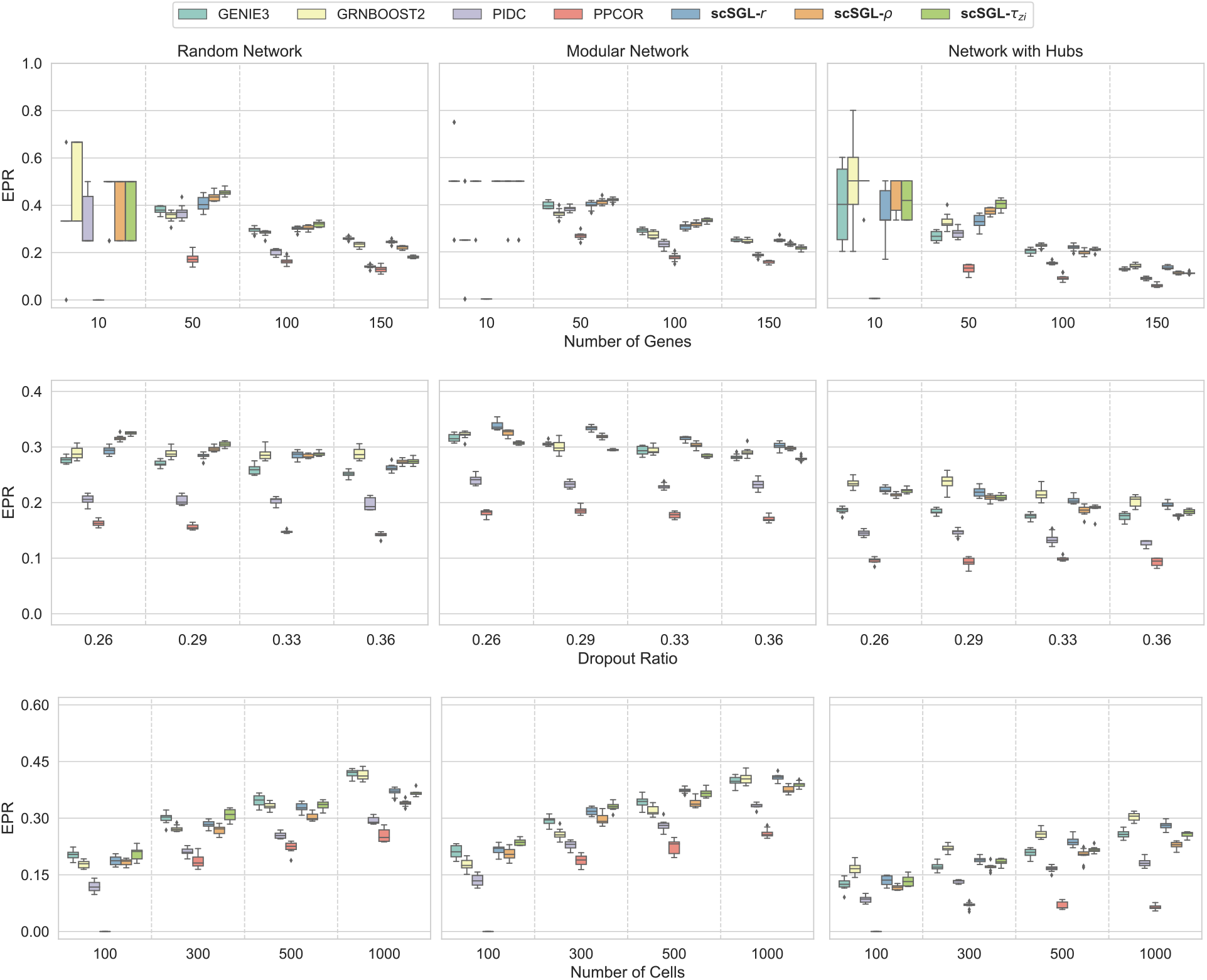
EPR values of various methods for synthetic datasets with three different topologies (random, modular and hub) and varying number of genes (top row), dropout ratio (middle row) and number of cells (bottom row).

**Figure 3:**
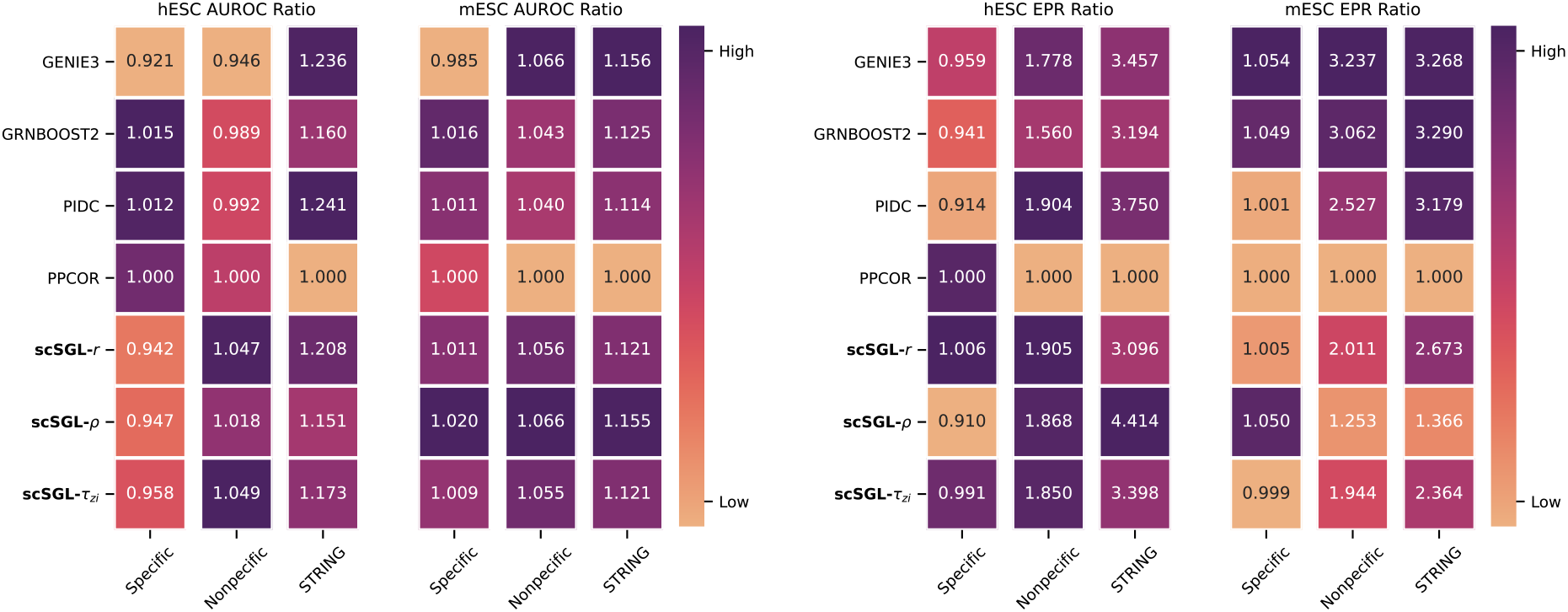
AUROC and EPR ratios of methods for two real-world scRNAseq datasets. Inferred graphs are compared to three different gene regulatory databases.

## Notes

### Competing Interest Statement

The authors have declared no competing interest.

### Summary of Updates

The proposed signed graph learning approach is extended to be able to use kernels. The kernelized version allows using different association metrics developed for gene expression. These metrics are used to reveal nonlinearity and handling dropouts. The results section is extended significantly.

